# Locations in the Neocortex: A Theory of Sensorimotor Object Recognition Using Cortical Grid Cells

**DOI:** 10.1101/436352

**Authors:** Marcus Lewis, Scott Purdy, Subutai Ahmad, Jeff Hawkins

## Abstract

The neocortex is capable of anticipating the sensory results of movement but the neural mechanisms are poorly understood. In the entorhinal cortex, grid cells represent the location of an animal in its environment, and this location is updated through movement and path integration. In this paper, we propose that sensory neocortex incorporates movement using grid cell-like neurons that represent the location of sensors on an object. We describe a two-layer neural network model that uses cortical grid cells and path integration to robustly learn and recognize objects through movement and predict sensory stimuli after movement. A layer of cells consisting of several grid cell-like modules represents a location in the reference frame of a specific object. Another layer of cells which processes sensory input receives this location input as context and uses it to encode the sensory input in the object’s reference frame. Sensory input causes the network to invoke previously learned locations that are consistent with the input, and motor input causes the network to update those locations. Simulations show that the model can learn hundreds of objects even when object features alone are insufficient for disambiguation. We discuss the relationship of the model to cortical circuitry and suggest that the reciprocal connections between layers 4 and 6 fit the requirements of the model. We propose that the subgranular layers of cortical columns employ grid cell-like mechanisms to represent object specific locations that are updated through movement.

## INTRODUCTION

Our brains learn about the outside world by processing our sensory inputs and movements. As we touch an object, survey a visual scene, or explore an environment, the brain receives a series of sensations and movements, a *sensorimotor sequence*.

Cortical areas that are traditionally viewed as sensory areas are known to integrate the motor stream into their processing. In vision, we perceive a stable image of the world, usually oblivious to the fact that our eyes are making multiple saccadic movements per second. As the eyes move, many neurons in the visual cortex that represent a particular stimulus anticipate the stimulus before it lands in the cell’s receptive field (Duhamel et al., 1992). In audition, responses in auditory cortex are predictively suppressed by motor signals (Schneider and Mooney, 2018). In somatosensation, when moving our fingers over familiar objects we quickly notice discrepancies suggesting we make tactile predictions that are specific to particular objects.

Predictive sensorimotor processing also occurs in the hippocampal formation. Grid cells (Hafting et al., 2005) and place cells (O’Keefe and Dostrovsky, 1971) represent an animal’s location, and they use a combination of sensory landmarks and self-motion cues to update their activity (Campbell et al., 2018). Another population of neurons selectively become active when an animal arrives at a location where a previously present object is missing (Tsao et al., 2013), indicating that the system is predictive.

Thus, different areas of the brain that seemingly play different roles in cognition display hallmarks of two common computations: integration of information over sensorimotor sequences, and prediction of sensory stimuli after movements. It is unclear how a network of neurons can extract reusable information from a sequence of sensations and movements, or how it can use this information to predict the sensory results of subsequent sequences. Simply memorizing sensorimotor sequences will lead to excessive learning requirements because for each sensed object there are many possible sensorimotor sequences.

Recent work from our lab (Hawkins et al., 2017) proposed that the neocortex processes a sensorimotor sequence by converting it into a sequence of *sensory features at object-centric locations*. The neocortex then learns and recognizes objects as sets of sensory features at locations that are in the reference frame of the object, and it predicts sensory input by referring to these learned object models. This approach integrates movement into object recognition, and forms representations of objects that generalize over novel sequences of movement. However in that paper we left open the neural mechanisms for computing such a location signal.

This paper extends our previous work by showing how the neocortex could represent and compute object-centric locations. Using this solution, we present a neural network model that learns and recognizes objects by processing sensorimotor sequences. We define a *sensor* to be the patch of skin or retina providing input to a particular patch of cortex. These patches of cortex can be thought of as cortical columns (Mountcastle, 1997). Drawing inspiration from how the hippocampal formation predicts sensory stimuli in environments, this model represents the sensor’s location relative to an object using an analog to grid cells, and it associates this location with sensory input. It then predicts sensory input by using motor signals to compute the next location of the sensor, and recalling the associated sensory feature. We propose that each patch of neocortex, processing input from a small sensory patch, contains all the circuitry needed to learn and recognize objects using sensation and movement. Information is also exchanged horizontally between patches, so movement isn’t always required for recognition (Hawkins et al., 2017), however this paper focuses on the computation that occurs within each individual patch of cortex.

There is a rich history of sensorimotor integration and learning internal models in the context of skilled motor behavior (Wolpert et al., 2011; Wolpert and Ghahramani, 2000). These have primarily focused on learning motor dynamics and kinematic control, such as reaching and grasping tasks. This paper focuses on a complementary problem, that of learning and representing external objects by integrating information over sensation and movement.

In the rest of this paper we first review the basic properties of grid cells in the entorhinal cortex. We then propose that the neocortex uses analogs of grid cells to model objects just as the hippocampal formation uses them to model environments. Building on the theoretical framework introduced in (Hawkins et al., 2017) we propose that every neocortical column contains a variant of this model. Based on cortical anatomy, we propose that cells in Layer 6 employ grid cell like mechanisms to represent object specific locations that are updated through movement. We propose that Layer 4 uses its input from Layer 6 to predict sensory input.

### How grid cells represent locations and movement

We first review how grid cells in the entorhinal cortex represent space and location. Although many details of grid cell function remain unknown, general consensus has emerged for a number of principles. Here we focus on two properties that are critical to our model: location coding and path integration.

Individual grid cells become active at multiple locations in an environment, typically in a repeating triangular lattice that resembles a grid (**Figure 1A**). The side length of these triangles in is known as the grid cell’s “scale”. A grid cell “module” is a set of grid cells that activate with the same scale and orientation but different positions, such that one or more grid cells will be active at any location (**Figure 1B**). If you sort the grid cells in a module by their relative firing locations, they form a rhombus-shaped tile. As the animal moves, a “bump” of activity moves across this rhombus (**Figure 1B and 1C**). The activity in a single module provides information on an animal’s location, but this information is ambiguous; many locations within the environment can lead to the same activity.

**Figure 1.**
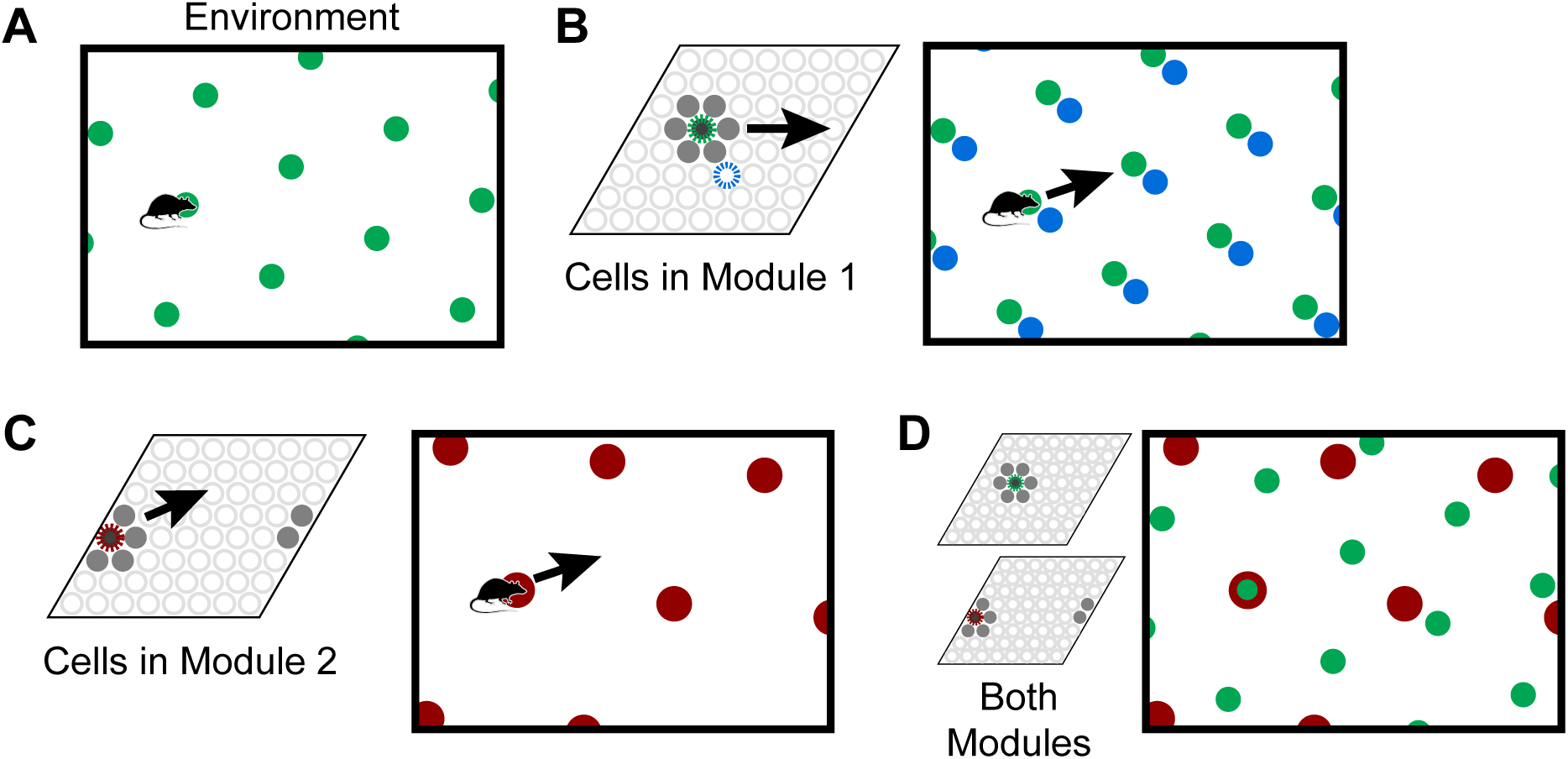
Grid cells represent locations in environments. **(A)** An individual grid cell becomes active at multiple locations (green circles) in an environment. The locations of activation form a repeating grid-like lattice. **(B)** A grid cell module (left) is a set of cells that share the same lattice scale and orientation but which activate at different relative positions in the environment. If you sort the cells by their relative firing locations, it forms a rhombus-shaped tile. As the animal moves, as shown by the arrow, a bump of cell activity will move in some direction through this rhombus. Two grid cells and their firing locations (green and blue) are highlighted. The grid cell module will activate cells at every location in an environment, but because of tiling, a single grid cell module cannot represent a location uniquely. **(C)** This figure shows how a second module tiles the same space differently. Each cell’s firing fields have a larger scale and a different orientation than the module in (A) and (B). The same movement of the animal as shown by the arrow causes the bump to move in a different direction and a different distance than the bump in the first module in (B). In this case, the bump overlaps the edge of the rhombus, so it wraps around. **(D)** Although a single module cannot represent locations in an environment uniquely, the activity across multiple modules can. Here we superimpose the firing patterns of the two modules. Note that when the green and red cells fire together, only one location is possible. The larger the number of modules, the more locations that can be represented uniquely.

To form a unique representation requires multiple grid cell modules with different scales or orientations (**Figure 1C and 1D**). For illustration purposes say we have 10 grid cell modules and each module can encode 25 possible locations via a bump of activity. These 10 bumps encode the current location of the animal. Notice, if the animal moves continuously in one direction the activity of individual modules will repeat due to the tiling, but the ensemble activity of ten modules is unlikely to repeat due to the differences in scale and orientation between the modules. The representational capacity formed by such a code is large. In our example the number of unique locations that can be represented is 25^10^ ≈ 10^14^. A review of the capacity and noise robustness of grid codes can be found in (Fiete et al., 2008; Sreenivasan and Fiete, 2011).

As an animal moves, the active grid cells in a module change to reflect the animal’s updated location. This change occurs even if the animal is in the dark (Hafting et al., 2005), telling us that grid cells are updated using information about the animal’s movement. This process, called “path integration”, has the desirable property that regardless of the path of movement, when the animal returns to the same physical location, then the same grid cells will be active (**Figure 2A**). Path integration is imprecise so in learned environments sensory landmarks are used to “anchor” the grid cells and prevent the accumulation of path integration errors (McNaughton et al., 2006; Ocko et al., 2018).

**Figure 2.**
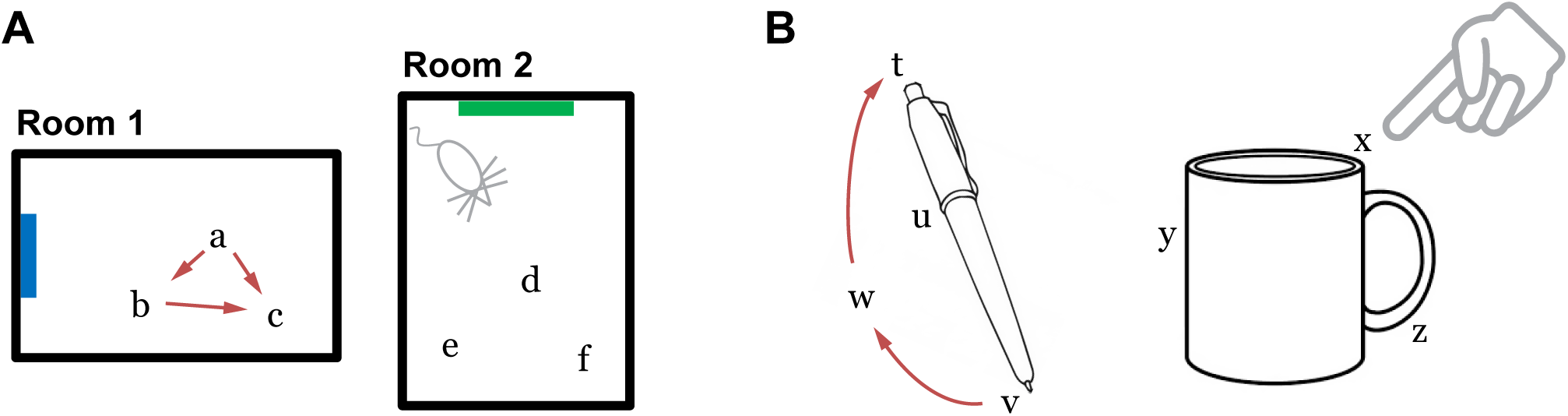
(A) Grid cells in the entorhinal cortex represent locations of a body in an environment. The location representations are updated by movement (Room 1). The path integration property ensures that the representation of location *c* is independent of the path taken to get there. Locations are unique to each environment such that the representations of locations *a, b*, and *c* are distinct from the representations of any point in Room 2. **(B)** We propose that the neocortex contains grid cell analogs that represent locations relative to an object. The location representations are unique to each object.

A final important property is that the location representations can be unique to each environment. If upon first entering a new environment each grid cell module activates a random bump to represent the current location. Then all the location representations that the animal can move to in that environment will, with high probability, be unique to that environment. The initial random starting point thus implicitly defines a unique location space for each environment, including locations that have not yet been explicitly visited. Since each module independently integrates motion information, path integration properties automatically hold for each new environment. Consequently, path integration can be learned once for each module and then reused across all environments. The location space for each new environment will be a tiny subset of the full space of possible cell activities (in our example above the full space contains 25^10^ points), thus the capacity for representing environments is quite large. When reentering a previously learned environment, learned associations between sensory cues and grid cells are used to “re-anchor” or reactivate the previous location space.

To summarize the above properties, a set of grid cell modules can unambiguously represent locations in an environment. These locations can be path integrated via movement, and environmental landmarks are used to correct path integration errors. By choosing random starting points within modules, unique location spaces can be defined for each environment. The space of all possible cell activations grows exponentially with the number of modules, thus the capacity for representing locations and environments is large.

## MODEL

We propose that grid cell equivalents exist throughout the neocortex. Rather than representing the location of the animal in an environment, we propose that cortical grid cells represent the location of sensory patches, for example the tip of a finger, in the reference frame of an object (**Figure 2B**). Similar to traditional grid cells, cortical grid cells define a unique location space around each object. As a sensor moves, populations of grid cells representing each sensory patch’s location will path integrate through unique location spaces. The relative locations of features on an object can thus be used as a powerful cue for disambiguating and recognizing objects.

Our network model integrates information over sensorimotor sequences, associating unique location spaces with objects and then identifying these location spaces. To outline the mechanism, let us first consider the question: how might a rat recognize a familiar environment? It must use its sensory input, but a single sensory observation may be insufficient to uniquely identify the environment. The rat thus needs to move and make multiple observations.

To combine information from multiple sensory observations, the rat could use each observation to recall the set of all locations associated with that feature. As it moves, it could then perform path integration to update each possible location. Subsequent sensory observations would be used to narrow down the set of locations and eventually disambiguate the location. At a high level, this general strategy underlies a set of localization algorithms from the field of robotics including Monte Carlo / particle filter localization, multi-hypothesis Kalman filters, and Markov localization (Thrun et al., 2005).

Our model uses this strategy to recognize objects with a moving sensor. The model uses each sensed feature to recall locations where it has previously sensed this feature, activating a superposition of these previously learned location representations. As the sensor moves, the network performs path integration on each of these recalled locations, i.e. the movement signal shifts the superposition of locations within each grid cell module. This updated location predicts a set of possible features, and the sensory input is used to confirm a subset of these predictions. Thus, with each subsequent sensation the network will narrow down this list of locations until it uniquely identifies a specific location on a specific object that is consistent with the sequence of sensations and movements.

Sensory features are known to invoke grid cell activity associated with familiar environments (Barry et al., 2007). Our model achieves this by learning associations between sensory input and the currently active grid cells at each location.

In our model, grid cell modules represent ambiguity by having multiple simultaneous bumps of cell activity, constituting a superposition of location representations. We refer to this set of simultaneously-active representations as a *union* of locations. The system is capable of path integrating unions of locations; during movement, every active bump in a module is shifted.

This localization algorithm assumes that the animal always knows the direction that it is moving in the reference frame of its environment. Determining this direction requires the animal to first perform *heading retrieval*, a computation that occurs somewhat independently of localization in the brain (Julian et al., 2018b). Our model doesn’t include an analog of heading retrieval. The model assumes it is given movement vectors in the reference frame of the object. In the discussion we briefly describe how this model could be extended to perform an analog of heading retrieval and hence build orientation-invariant models of objects.

### Model description

Our two-layer model consists of two populations of neurons and four primary sets of connections **(Figure 3)**. Later in the “Mapping to biology” section we propose which cortical populations implement this circuit. For each movement of the sensor, the network goes through a progression of stages, processing the motor input followed by the sensory input. Each stage corresponds to using the connections from one of the numbered arrows in **Figure 3**. We show an example of the network going through these stages three times in **Figure 4**.

**Figure 3.**
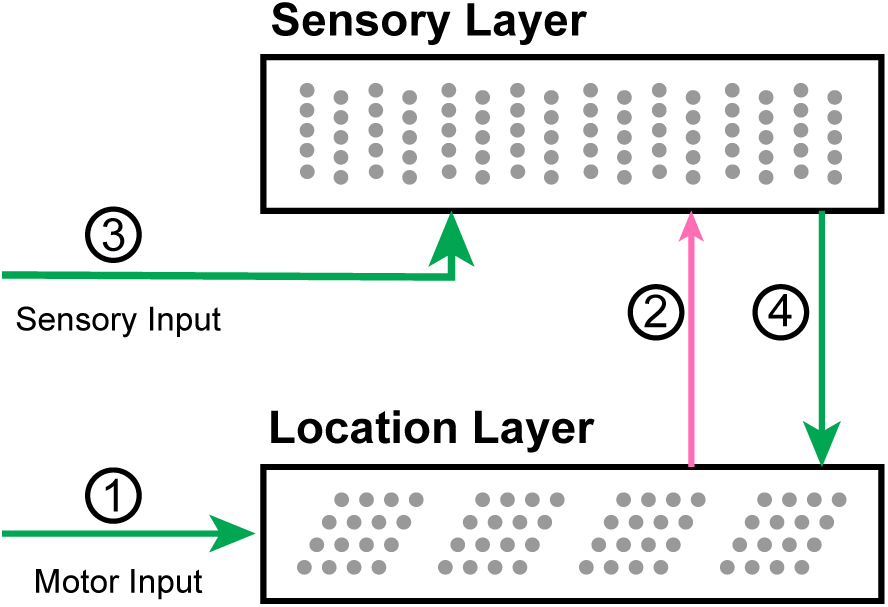
A diagram of the network with arrows indicating the main connections. (1) Motor input shifts the activity in the location layer, which consists of a set of independent grid cell modules. (2) The active location cells provide modulatory predictive input to the sensory layer. (3) Sensory input activates cells in the sensory layer. (4) The location is updated by the new sensory representation.

**Figure 4.**
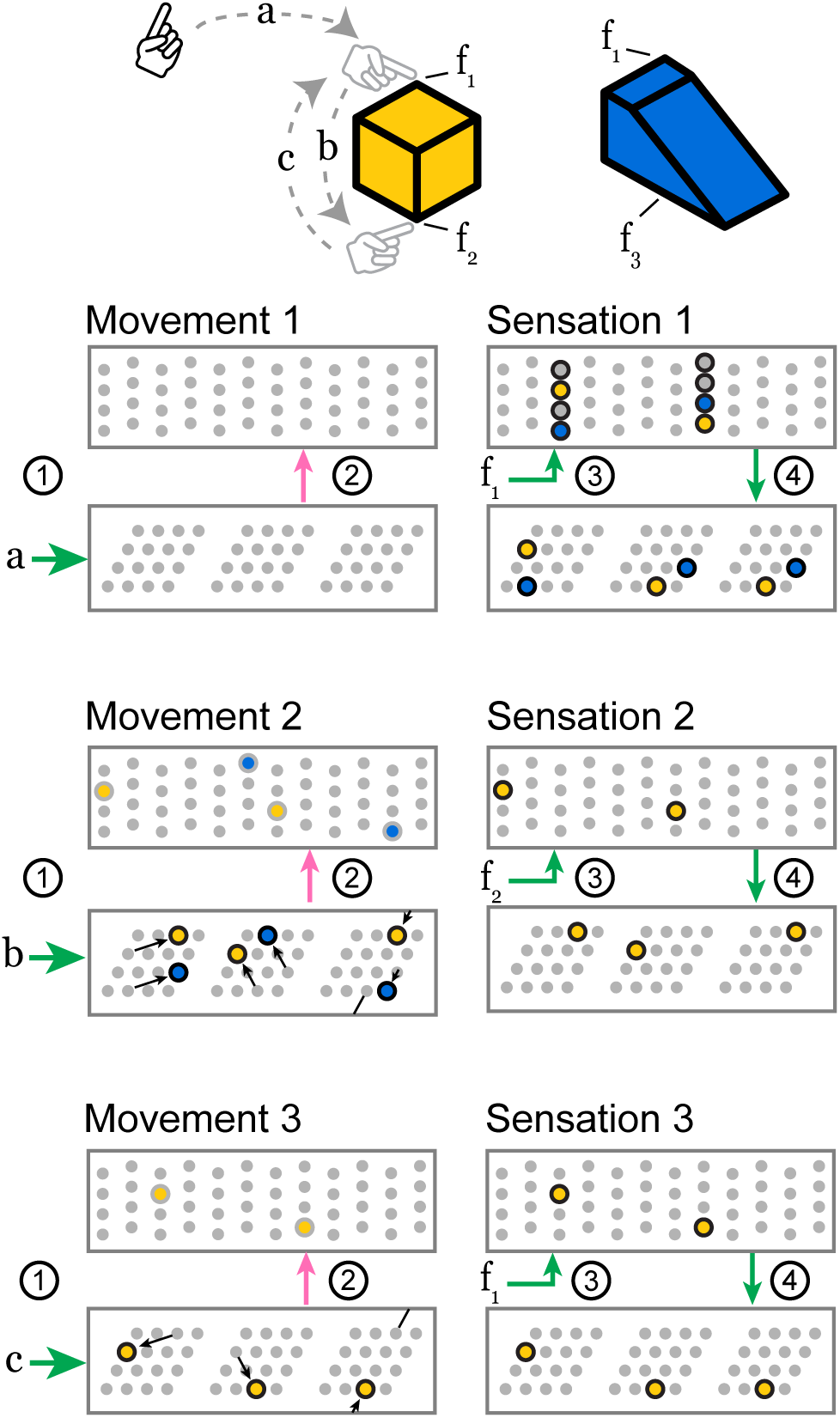
As the sensor moves over a previously learned object, these two layers receive a sensorimotor sequence and recognize the object. Features *f*_1_, *f*_2_ and *f*_3_ indicate the sensory input invoked by touching the objects at the indicated locations. Motor commands *a, b*, and *c* indicate the motor input received by the network when the sensor makes a movement. The objects are colored to relate them to active cells below. We show three movements, each consisting of the four stages above, and we draw snapshots of the network at the end of stages 2 and 4. The stages 1 through 4 correspond to the connections in Figure 3. **Movement 1.** The network receives a movement command, and nothing happens because it doesn’t have a current location representation. **Sensation 1.** The sensor senses feature *f*_1_ which provides input to every cell in a set of minicolumns. None of the cells were predicted, so all become active. This feature has been learned on two objects, so this set of active cells contains two feature-at-location representations, shown in yellow and blue. These representations drive a pair of location representations to become active. This union of activity encodes the two possible locations that could have produced the sensation. **Movement 2.** Motor input *b* causes each module to perform path integration, shifting its bump according to its scale and orientation. The newly active cells provide modulatory input to the sensory layer which predicts two different potential features, *f*_2_ and *f*_3_, priming two representations to become active. **Sensation 2.** The sensor senses feature *f*_2_, and only the predicted cells in the *f*_2_ mini-columns become active. The other predicted cells do not activate. This active representation drives a single representation to become active in the location layer. At this point, the network has identified the cube. **Movement 3 and Sensation 3.** Subsequent movements maintain the unambiguous representation as long as the sensed features match those predicted by the path-integrated locations. A movement back to the original location, for instance, causes a prediction only for the *f*_1_ representation specific to that location on the cube.

**Stage 1.** Motor input arrives before the sensory input and is processed by the location layer, which consists of grid cell modules. If this layer has an active location representation, it uses the motor input to shift the activity in each module, computing the sensor’s new location.

**Stage 2.** This updated grid cell activity propagates to the sensory layer and causes a set of predictions in that layer.

**Stage 3.** The sensory layer receives the actual sensory input. The predictions are combined with sensory input. The new activity is a union of highly sparse codes. Each sparse code represents a single sensory feature at a specific location that is consistent with the input so far.

**Stage 4.** The sensory layer activity propagates to the location layer. Each module activates a union of grid cells based on the sensory representation. The location layer will contain a union of sparse location representations that are consistent with the input so far.

After the fourth stage the next motor action is initiated and the cycle repeats. The next few sections describe the network structure and each of these 4 stages in detail, as well as the learning process.

### Network structure

We compute the network activity through a set of discrete timesteps. Each time step *t* consists of a progression of the 4 stages outlined above. Here we describe the neuron model and network structure before describing the stages in detail.

Each neuron in the network is a discrete time neuron with multiple independent dendritic segments. 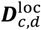 and 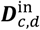 each denote a vector which specifies the synapses of dendrite *d* on cell *c* in the location layer and sensory layer, respectively. Each neuron has a binary output, and synapse weights are either 0 or 1. The number of synapses on each dendrite after learning is generally small and as a result these vectors are highly sparse. The neuron model with independent dendritic segments is closely related to the structure of the Poirazi-Mel neuron (Poirazi and Mel, 2001) and the existing experimental literature on active dendrites in pyramidal neurons (Antic et al., 2010; Major et al., 2013). Multiple dendritic segments enable each neuron to robustly recognize independent sparse patterns, and thus be associated with multiple location or sensory contexts. Although the activity of a single neuron can be ambiguous, we have shown in (Ahmad and Hawkins, 2016; Hawkins and Ahmad, 2016) that the activity of a network of such neurons can represent sparse distributed codes that are highly unique to specific contexts. In addition, with a sufficiently large number of cells, sparse representations enable such networks to represent a union of patterns (Ahmad and Hawkins, 2016) with a low probability of false match errors (up to a limit).

### Sensory and location layers

The network presented here is an extension of the work in (Hawkins et al., 2017), and the sensory layer is identical in structure to the sensory input layer in that paper. As in that paper, the layer is organized into a set of mini-columns (Buxhoeveden, 2002) such that all cells within a mini-column have identical feedforward receptive fields but inhibit each other. Dendritic segments of the neurons in the sensory layer have a modulatory effect. An active segment does not by itself cause the cell to become active. When a cell with an active dendritic segment recognizes feedforward input, that cell will inhibit any other cells within the mini-column that do not have active segments (see Stage 3 below). The active cells of mini-column *i* are denoted by the binary array 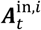 The layer activity 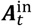 consists of the concatenation of all of the mini-column activities 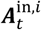.

The location layer is structured as a set of grid cell modules, each containing the same number of neurons. The active neurons of module *i* at time *t* are denoted by the binary array 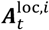, and the layer activity 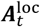 consists of the concatenation of all of the module activities 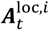. Dendritic segments of the neurons in this layer have a driving effect; a cell will become active if any of its dendritic segments become active. During inference, activity in the location layer is updated once in response to movement and then again in response to sensory-derived input. We denote these two activation states by the vectors 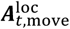, and 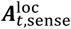.

The location layer projects to the dendritic segments of the sensory layer (**Figure 3**, connection 2). Thus 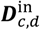 is a vector with the same length as 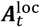 where a 1 represents a connection to a cell in the location layer. The sensory layer projects to the dendritic segments of the location layer (**Figure 3**, connection 4). Thus 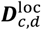 is a vector with the same length as 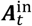 where a 1 represents a connection to a cell in the sensory layer. These vectors are generally extremely sparse as they connect to sparsely active cells during learning (see section on learning below).

In each timestep, dendritic segments that receive sufficient input become active. The binary vectors 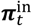 and 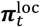 denote cells in each layer that contain at least one active dendritic segment. These denote whether each cell was *predicted* from the other layer’s activity. Designating *θ*^in^ and *θ*^loc^ as dendritic thresholds,

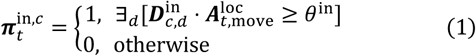

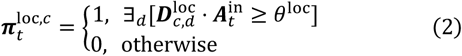

### Stage 1: Use movement to update location layer

The location layer consists of a set of independent grid cell modules, each with its own scale and orientation. We follow notation and assumptions of other grid cell models and analyses (Ocko et al., 2018; Sreenivasan and Fiete, 2011). Within a module, the active cells are always part of a Gaussian bump of cell activity centered at a position, or *phase*, within the module’s tile (**Figure 1B**). We designate the phase of module *i*’s bump as 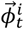. Each cell within a module is centered at a phase, and its activity at time *t* is proportional to its nearness to 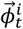. The network responds to movement commands by updating 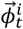 for each module.

Movement will shift each module’s bump according to the module’s scale and orientation. A 2D transform matrix ***M***_*i*_ associated with each module represents how the module converts a movement vector for the sensor into a movement vector for the bump. Denoting the scale of module *i* as *s*_*i*_ and the orientation as *θ*_*i*_, this transformation matrix is:

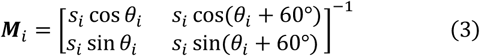

This operation converts the movement vector into a 60° basis, arranging each grid cell’s firing fields into a triangular rather than square lattice. It also scales and rotates the movement vector to set the scale and orientation of the lattice.

Each module receives the same 2D movement vector 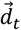, and it shifts its bump as follows:

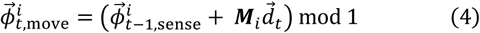

The modulo operation confines each bump of activity to have a phase within the 2D range [0, 1) × [0, 1). In the sorted view of a grid cell module (**Figure 1B**), the phase [0, 0] corresponds to the lower left corner while [0.9, 0.9] corresponds to the upper right corner. The bump of cell activity is centered at this phase, and we model the bump as having a Gaussian shape to qualitatively match the shape of the firing fields of observed grid cell responses (Monaco and Abbott, 2011).

In addition to these properties, our model requires a grid cell module to be capable of path integrating multiple bumps simultaneously. Each module represents uncertainty by activating multiple bumps, one for each possible location. We refer to this as a *union* of locations. We designate a module’s set of bumps as 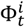. With every movement, we apply Eq. (4) to every phase in 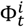.

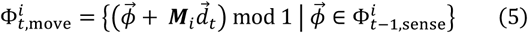

Rather than simulating individual cell dynamics explicitly, each module in the location layer simply maintains a list of activity bump phases and updates each one according to Eq. (5). The location layer then outputs the binary vector 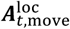 by thresholding the activity of each cell within the module (see Model Details). In the discussion we review models of individual grid cell dynamics and discuss their compatibility with unions.

### Stage 2: Use updated location to form sensory predictions

In Stage 2, we compute which sensory features are predicted by the location layer. In our network these predictions are represented by 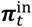, the activity of distal dendritic segments of cells in the sensory layer. We compute 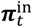 from 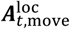 using Eq. (1) above.

Each sensory feature that has been encountered at any of the current possible locations will be predicted. Note that because 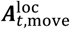 is a concatenation of all grid cell modules, the predictions are based on highly specific location codes.

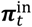 has a modulatory effect on the activity of the sensory layer, as described in Stage 3.

### Stage 3: Calculate activity in sensory layer

In Stage 3, the sensory layer uses the sensed feature to confirm correct predictions and to disregard incorrect predictions. When no predictions are correct, it activates all possible feature-at-location representations for the feature. This stage is responsible for both activating and narrowing unions.

The sensory layer is identical to the sensory input layer in (Hawkins et al., 2017). In this layer, all the cells in a mini-column share the same feedforward receptive fields. Each sensory feature is represented by a sparse subset of the mini-columns, denoted by 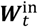. When a cell is predicted, it is primed to become active, and if it receives sensory input it will quickly activate and inhibit other cells in the mini-column.

The active cells within the sensory layer are selected by considering 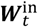 and by considering which cells are predicted by the location layer, i.e. which cells have an active dendritic segment. If a cell is predicted and it is in a mini-column in 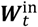, it becomes active. If a mini-column in 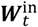 has no predicted cells, every cell in that mini-column becomes active.

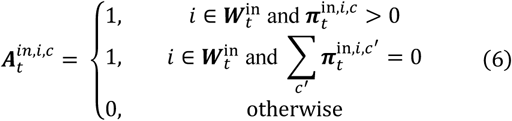

If the location layer has uniquely identified the current location, there will be exactly one active cell in each mini-column in 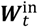 encoding this sensory feature at the current location. If the location layer contains a union of locations, the sensory layer will represent this feature at a union of locations. Note that the location layer may predict features that are not represented in 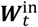 These mini-columns will not contain any activity after this step (**Figure 4**, Sensation 2) and the set of possible objects is thus narrowed down.

### Stage 4: Update location layer based on sensory cues

In Stage 4, the activity in the sensory layer recalls locations in the location layer. 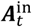 is used to compute 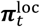 (Eq. (2)) which drives activity in the location layer. The list of activity bump phases in each module is replaced by a new set of bumps driven by sensory input. For each cell in the location layer that has a corresponding active dendritic segment, the module activates a bump centered on that cell.

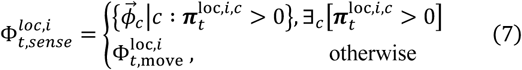

This new set of bumps is often very similar to the previous set, as in **Figure 4,** Sensation 3. Note that this step happens during inference only. During learning, the location layer doesn’t update in response to sensory input; we simply assign 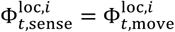.

The active cells in the location layer 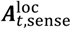 are computed from 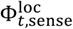 (see Model Details). Note that since connections from the location layer to the sensory layer cannot drive activity, recurrence in the network is minimal (see Model Details).

### Learning

In our model the learning process involves associating locations with sensory features, and vice versa. These associations are learned on the distal dendritic segments, as specified in Eq. (8) and Eq. (9). Learning in this model always consists of the active cells in the two layers forming reciprocal connections. At the start of training on a new object, each module in the location layer activates a bump at a random phase. This instantiates a random location space that is specific to that object. For the rest of training, we provide a sequence of motor and sensory inputs, calculating 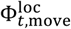 as before, shifting each module’s bump with each movement.

Each sensory input is represented by a set of mini-columns 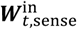. Following Eq. (6) above, if this part of the object hasn’t been learned yet there will be no predictions in the sensory layer and every cell in these mini-columns will become active. In this case a random cell in each active mini-column is selected as the cell to learn on, i.e. to represent this sensory input at this location. If this part of the object has been learned, there will be predictions in the sensory layer. In this case the cells corresponding to the existing active segments are selected to learn on. 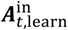 represents these learning cells for the current time step.

Each active cell in 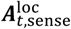 and 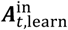 selects one of its dendritic segments *d*′ and forms connections between this segment and each active cell in the other layer.

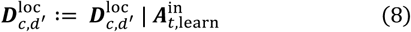

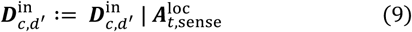

Here we designate “|” as bitwise OR, implying that the existing connections on the dendritic segment are unaffected when these new connections are formed.

## SIMULATION RESULTS

We ran simulations to test the ability of our model to learn the structure of objects and accurately predict sensory features from motor sequences. Our model is agnostic to the specific feature extraction technique, thus we focus on testing the set of neural computations that occur after features have been extracted. Thus, these experiments tested this set of neural computations and weren’t intended to be a general benchmark for object recognition. Our goal was to understand whether the model can accumulate information over sensorimotor sequences and to understand when the model reaches its breaking point.

In the simulations, the model’s input consisted of sequences of sensory features and movements. The model’s task was to learn an object from a single sensorimotor sequence and then recognize that object from a different sensorimotor sequence. In **Figure 5** we show an illustration of our test objects and the input sequences. Each object consisted of ten points chosen randomly from a four-by-four grid. We placed a feature at each point, choosing each feature randomly with replacement from a fixed feature pool. Features were shared across objects and a given feature could occur at multiple points on the same object.

**Figure 5.**
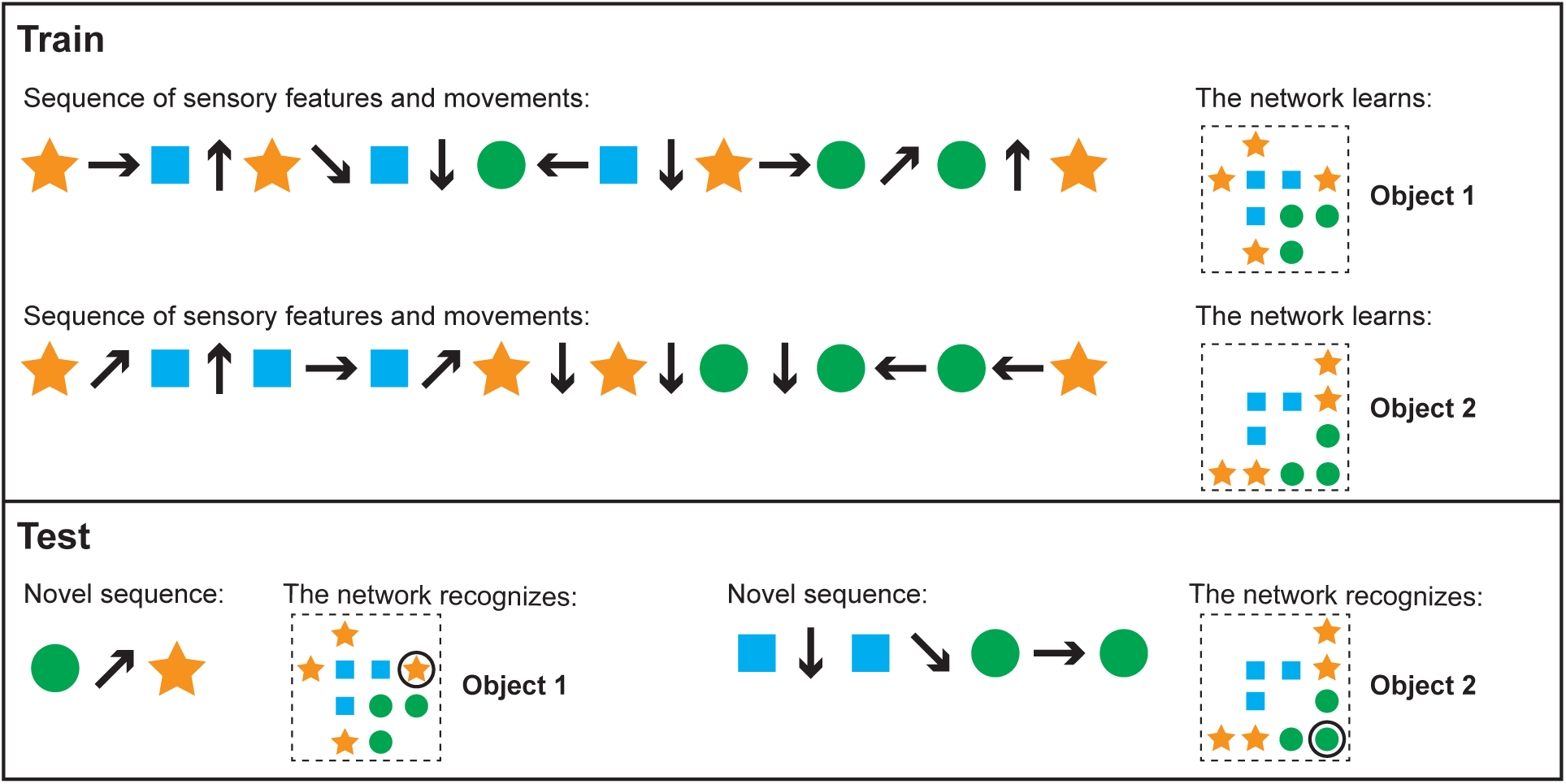
Our model learns and recognizes objects from sensorimotor sequences. The order of movements and sensations is not critical for either learning or recognition. We ran simulations to test these properties. This cartoon figure illustrates a training and test run of one of these simulations. **(Top)** Two objects are shown (right side). Each object is composed of the same sensory features (circle, square, star) in different spatial arrangements. The network learns the objects via a sequence of sensory features and movements (left). Note that the model learns the relative arrangement of the features (right), not the sensorimotor sequence itself (left). **(Bottom)** The network can recognize objects with novel sensorimotor sequences. The left input sequence consists of a sensed circle, a diagonally upward movement, and a sensed star. Although the network has never experienced this particular sensorimotor sequence before, it detects that this sequence matches a portion of object 1 and doesn’t match any other objects. In this example, one movement and two sensations is sufficient to recognize the object and the sensor’s current location on that object. The right sensorimotor sequence matches locations on both objects until the fourth sensory input, so it takes three movements and four sensations to recognize that this is object 2.

For every simulation, we trained the network by visiting each point on each object once. For each point, we stored the activity in the location layer in a separate classifier. We then tested the network on each object by traversing each of the object points in random order. Importantly, the network received a motor input for each of these random transitions between features, enabling it to properly accumulate information over time. As the sensor traversed the object, we tested whether the location representation exactly matched the classifier’s stored representation for that point on that object. If it matched this representation and if this representation was unique to this object, we considered the object to be recognized. If the network never converged to a single location representation after four complete passes over the object, or if it ever converged on a wrong location, we considered this a recognition failure. (In these experiments, the model always either converged to a single location or didn’t converge at all.) It is possible for the network to converge on the correct location representation even if that location isn’t unique to the object, for example if there aren’t enough modules to create a unique code. In the sections ahead, we specifically note when this occurred.

We set the sensory layer to have 150 mini-columns and 16 cells per mini-column. Each sensory feature activated a predetermined, randomly selected set of 10 mini-columns which comprised the 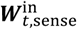 of timestep *t*. We varied the number of cells per module and the number of modules in the location layer. For these simulations we varied the module orientations but not the scales. The orientations were evenly spaced along the 60° range of possible orientations. Each module used the same scale, and this scale was fixed to be half the maximum width of the objects. We set the dendritic thresholds *θ*^loc^ and *θ*^in^ to 8 and ⌈*n* * 0.8⌉, respectively, where *n* is the number of modules in the location layer and ⌈ ⌉ is the ceiling operator.

### Cell activity converges to a unique location representation

We begin by demonstrating the cell activity in a typical recognition task, and we show how it varies as the network learns more objects. In **Figure 6** we show the actual grid cell activity for several modules in the location layer as the network sensed different objects. The network was first trained on 50 objects, each with ten features drawn from a pool of 40 features. The activity changes with each new sensation, first via path integration shifts based on the movement, followed by narrowing of the activity to only the locations consistent with the newly sensed feature. The network can take a different number of sensations to narrow to a single representation per module. Once the network has narrowed, the activity in a single module may be ambiguous but the set of active cells across all modules (only three of ten modules are shown) uniquely encode the object and location.

**Figure 6.**
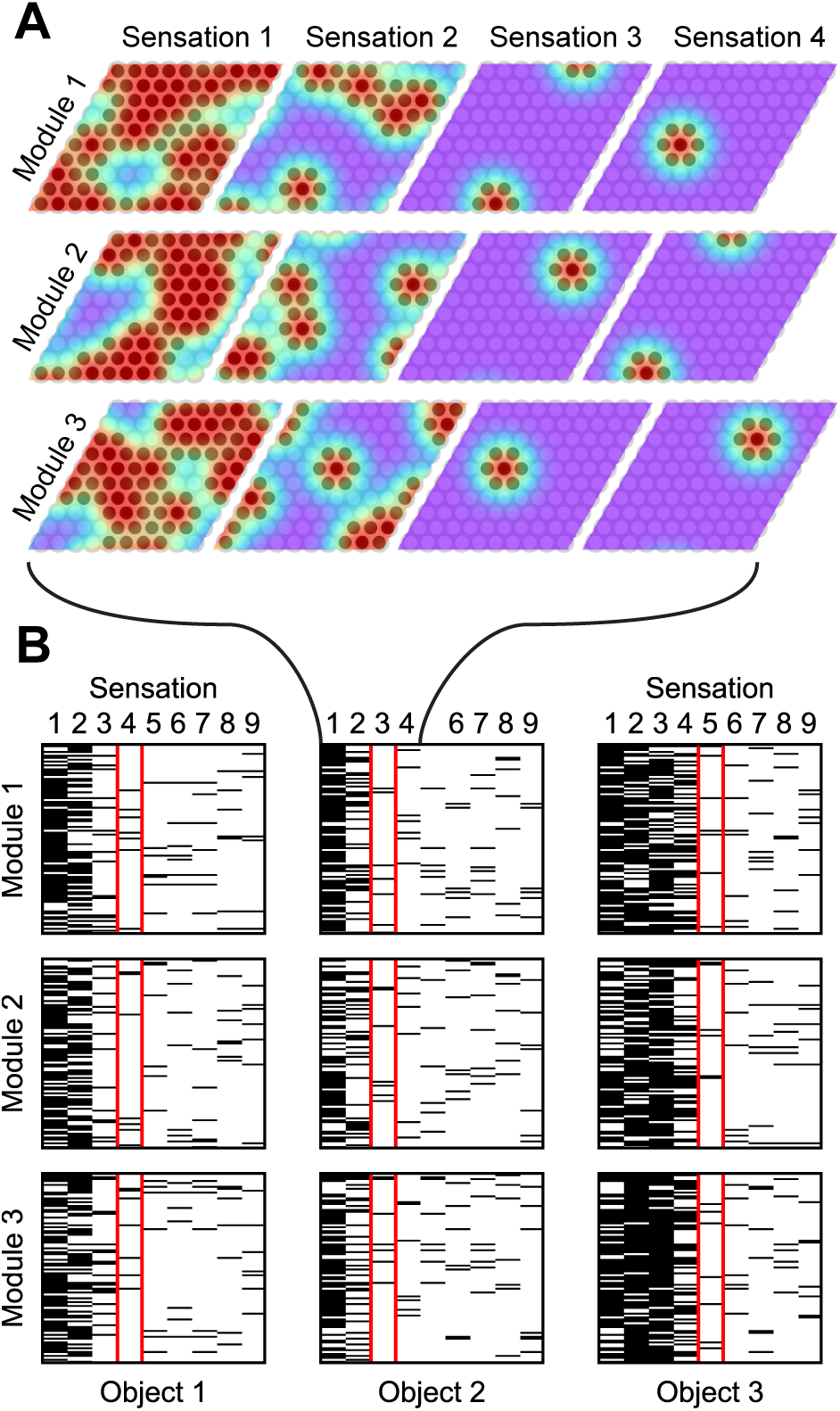
As the network recognizes an object, it converges onto a sparse activation. **(A)** Location layer cell activity in three (out of ten) grid cell modules while recognizing a single object. The bumps of activity are shown in color; red indicates a high firing rate. The location representation, shown as darkened circles, consists of cells that are sufficiently active. Each movement shifts the activity and each sensation narrows the activity to the cells that predict the new sensation. Cell activity converges onto a sparse representation by Sensation 3 and remains sparse afterward. **(B)** Cell activity in the same three modules as (A), shown for additional sensations and for two additional objects. Each module has 100 cells. The black lines indicate that the cell is active during the indicated sensation. After the first sensation, the location codes are very ambiguous in all cases. Depending on how common the sensed features are, the module activity narrows at different rates for the different objects. The sensation highlighted in red shows the first step in which the object is unambiguously determined. From this point on, the module activity shifts with each movement but remains unique to the object being sensed. (These simulations used a unique feature pool of size 40.)

The recognition process always follows this template. The initial sensory input typically causes dense activation, assuming the sensory feature is not unique to a single location. With subsequent movement and sensation this activity becomes sparser and eventually converges on a single representation. In **Figure 7A** we aggregate the cell activity from **Figure 6** across all objects and all modules to show the average cell activation density after each sensation. As the network learns more objects, the initial density and the convergence time increase because the network recalls more locations-on-objects for each sensory input, and it has to disambiguate between more objects.

**Figure 7.**
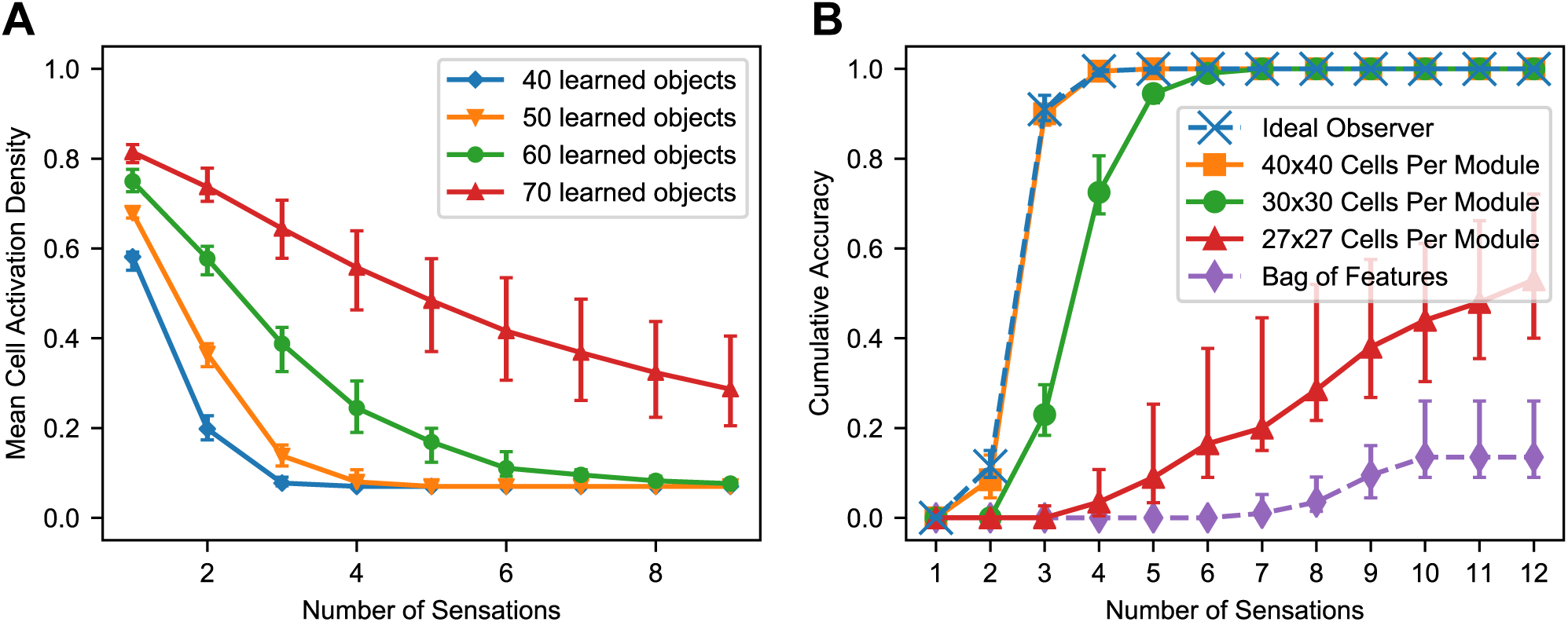
**(A)** With multiple sensations, the location layer activity converges to a sparse representation. Using the same simulation from Figure 6, we show the activation density after each sensation, averaged across all objects and modules. With additional sensations, the representation becomes sparser until the object is unambiguously recognized. With more learned objects, the network takes longer to disambiguate. The activation density of the first sensation increases with the number of learned objects. If the initial activation density is low, the network converges very quickly. If it’s high, convergence can take longer. **(B)** Comparison of this network’s performance with the ideal observer and a bag-of-features detector. Each model learned 100 objects from a unique feature pool of size 10. We show the percentage of objects that have been uniquely identified after each sensation, averaged across all objects. The ideal model compares the sequence of input features and relative locations to all learned objects while the bag-of-features model ignores relative locations. With a sufficiently large number of cells per module, the proposed neural algorithm gets similar results to the ideal computational algorithm. **(Both)** In both of these charts we repeat the experiment with 10 different sets of random objects and we plot the 5th, 50th, and 95th percentiles of each data point across trials.

### The model approximates an ideal observer

Ideally, the network should recognize an object as soon as it receives a sensorimotor sequence that uniquely matches that object. In **Figure 7B** we compare the recognition time with such an ideal observer. The ideal detector stores both the features and their relative locations during training and exhaustively checks the current sensorimotor sequence against the stored objects during testing. It yields a correct classification as soon as the object is unambiguous based on the features and relative locations sensed up to that point. We also include a bag of features model. It ignores location information and yields a correct classification if it can unambiguously determine the object based solely on the sensory features.

This network uses 10 modules, and the dataset contains 100 objects with 10 unique features. Very few objects can possibly be determined by a single sensation, but three or four sensations are sufficient. As we increase the number of cells in each module, the model’s performance approaches that of the ideal detector. With 40×40 cells per module, the two are near identical. The bag of features model often cannot uniquely identify the objects because many objects are different arrangements of identical sets of features.

### Near the model’s capacity limits, recognition time degrades

As this model learns more objects, it eventually begins taking longer to recognize objects than the ideal observer. If it’s pushed further, it eventually begins failing to recognize objects. In **Figure 7B**, the network with 30×30 cells per module requires more sensations to narrow down the object than the ideal observer, but it always recognizes the object eventually. The network with 27×27 cells per module often never recognizes the object after many sensations, indicating the network has been pushed beyond its capacity.

To understand the way that the system reaches a capacity limit, consider the density of activation in the location modules when a single feature is sensed. If a single feature occurs in many locations, then during learning the location layer and sensory layer reciprocally associate many cells in each module with that feature. Sensing that feature will cause a large percentage of the cells in the location layer to activate. If this percentage is too high, location representations that aren’t supposed to be active will be largely contained in this dense activation, resulting in false positives. In the worst case, the location layer will fail to extract anything useful out of this sensory input, because it will activate nearly every location representation as part of its dense activation. We found that the main influence on the model’s recognition time is whether the model is approaching its capacity limit. In the following section we characterize this capacity limit.

### Capacity varies with size of location layer, statistics of objects

The previous simulations investigated the time to converge onto a unique location. Here we consider the capacity of the network independent of convergence time. We compute the fraction of objects correctly classified after many sensations, and we define the capacity as the maximum number of objects the network can store while maintaining a 90% accuracy rate. While we use 10 locations per object for these simulations, the total number of locations (number of objects times the number of locations per object) is what matters.

It is worth noting that this definition of capacity is a measure of the model’s ability to recognize objects, not simply a measure of its ability to store objects. The network could potentially store many more predictive models of objects, but after some point it will be unable to recognize those objects from sensory input via this model’s circuitry. The model has multiple potential bottlenecks that could determine this breaking point: the representational capacity for sensory features, the representational capacity for locations, the number of patterns a cell can learn via independent dendritic segments, and the network’s ability to represent multiple locations simultaneously. By changing the model parameters and input data in different ways, any of these could become the bottleneck, but we found that the most unavoidable of these potential bottlenecks was the last one.

Because our experiments focused on scenarios in which the same features occur on multiple objects, the model begins running into capacity limits when the sensory input causes the location layer to activate a union of representations that is large enough to cause false positives in the sensory layer. This occurs when the union contains large portions of location representations that aren’t supposed to be active. The likelihood of this event is influenced by the model’s number of modules, the number of cells per module, and the statistics of objects. In **Figure 8**, we characterize the impact of these variables on the model’s capacity.

**Figure 8.**
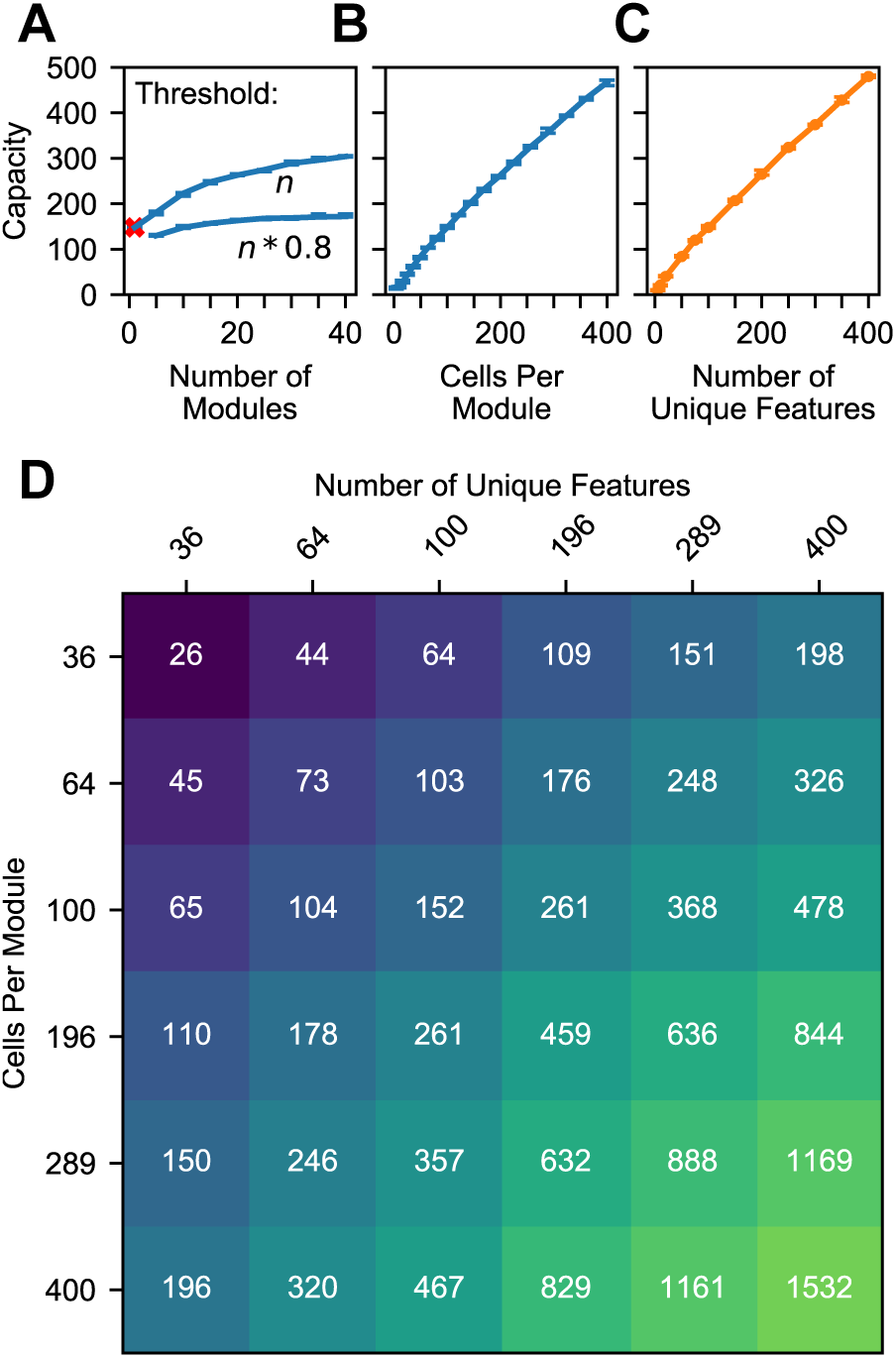
**(A-C)** Model capacity changes with model parameters (blue) and object parameters (orange). The network’s capacity is the maximum number of objects that it can learn while maintaining the ability to uniquely identify at least 90% of the objects. We repeat the experiment with 10 different sets of random objects and we plot the 5th and 95th percentiles of each capacity. **(A)** Increasing the number of modules in the network increases capacity sublinearly, and this impact depends on the dendritic threshold of the neurons reading this location representation. The two lines show two different thresholds relative to *n*, the number of modules. With only one module the network can successfully converge to one location, but that location isn’t object-specific (red ‘x’). **(B)** Increasing the number of cells increases capacity linearly. **(C)** Similarly, increasing the feature pool size also increases capacity linearly. **(D)** A heatmap showing the capacity of the network while varying the feature pool size and location module size. An increase in the number of cells per module can compensate for a decrease in the number of unique features.

The model’s capacity increases with the number of modules, though with diminishing returns. In **Figure 8A** we plot two lines to dissect the relationship between the number of modules and capacity. The top line shows the impact of having multiple modules, then the bottom line shows how this effect is reduced by our model’s lower dendritic thresholds. Even with a single module, the network is able to converge onto one location representation for a considerable number of learned objects. However, this location representation isn’t unique to the object, so it doesn’t qualify as recognizing the object; we denote this with a red ‘x’. Adding a few more modules ensures unique locations, and each additional module helps the model deal with large unions. They guard the model from having too many false positives when a large percentage of cells are active. More precisely, for each representation, the percentage of the cells that will be active due to randomness will be approximately equal to the activation density, with some variance, and having more modules reduces this variance. If the threshold is 100%, then the model can increase its capacity indefinitely by adding modules. However, to avoid depending on every neuron reliably firing, this model doesn’t use such a high threshold. In our model, neurons will detect a location if 80% of its cells are active. With this lower threshold the benefit of additional modules asymptotes, and no number of modules will be able to handle more than 80% activation density.

Previous analysis of grid cell codes (Fiete et al., 2008) showed that the code’s representational capacity increases exponentially with the number of modules. This holds true for this model, but this model’s bottleneck is not the size of its unique location spaces. This model’s performance depends on its ability to unambiguously represent multiple locations simultaneously. A grid cell code’s *union* capacity doesn’t scale exponentially with number of modules.

The model’s capacity increases linearly with number of cells per module **(Figure 8B)**. As we add additional cells, the size of a bump remains constant relative to the cells, so the bump shrinks relative to the module. Because each bump activates a smaller percentage of the cells in a module, a module can activate more bumps before the network reaches the density at which object classification starts failing.

The model’s capacity increases linearly with the number of unique features **(Figure 8C)**. This happens because it reduces the expected total number of occurrences of each feature, and hence the number of elements in each union. This indicates that the statistics of the world influence the capacity of the model, and it also means that this network can improve its capacity by adjusting the “features” that it extracts from sensory input.

Because these two latter parameters have independent linear relationships with capacity, they can compensate for each other. In **Figure 8D** we plot object capacities in a network with 10 modules. We show that increasing (decreasing) the number of cells per module and decreasing (increasing) the number of unique features by the same factor causes the capacity to remain approximately constant. This is illustrated by the approximate symmetry across the chart’s diagonal.

### The model recognizes an object if the object has at least one sufficiently uncommon feature

Up to this point, we’ve characterized this model’s ability to recognize objects by stating, “The network will reliably recognize an object if the network has learned fewer than *c* total objects,” and we’ve measured *c*. This characterization is built on many assumptions about objects. It’s desirable to be able to characterize the model in a way that isn’t specific to these assumptions.

Given that the model’s ability to recognize objects depends on the density of cell activity invoked by sensory features, we found we could characterize the model’s performance more directly by answering, “The network will reliably recognize an object if the object contains a feature with fewer than *k* total occurrences across all learned objects,” and measuring *k*. In another set of experiments (**Figure S1**), we generated objects using multiple alternate distributions of features. We found that the network’s breaking point relative to *c* (the number of learned objects) did indeed vary widely with the choice of feature distribution, while its breaking point relative to *k* (the number of locations recalled by sensing a feature) was much more consistent across distributions. This suggests that the network’s performance relative to *k* will hold true with real-world statistics. Using both of these metrics, *c* and *k*, we summarize all of these results in **Figure 9**.

**Figure 9.**
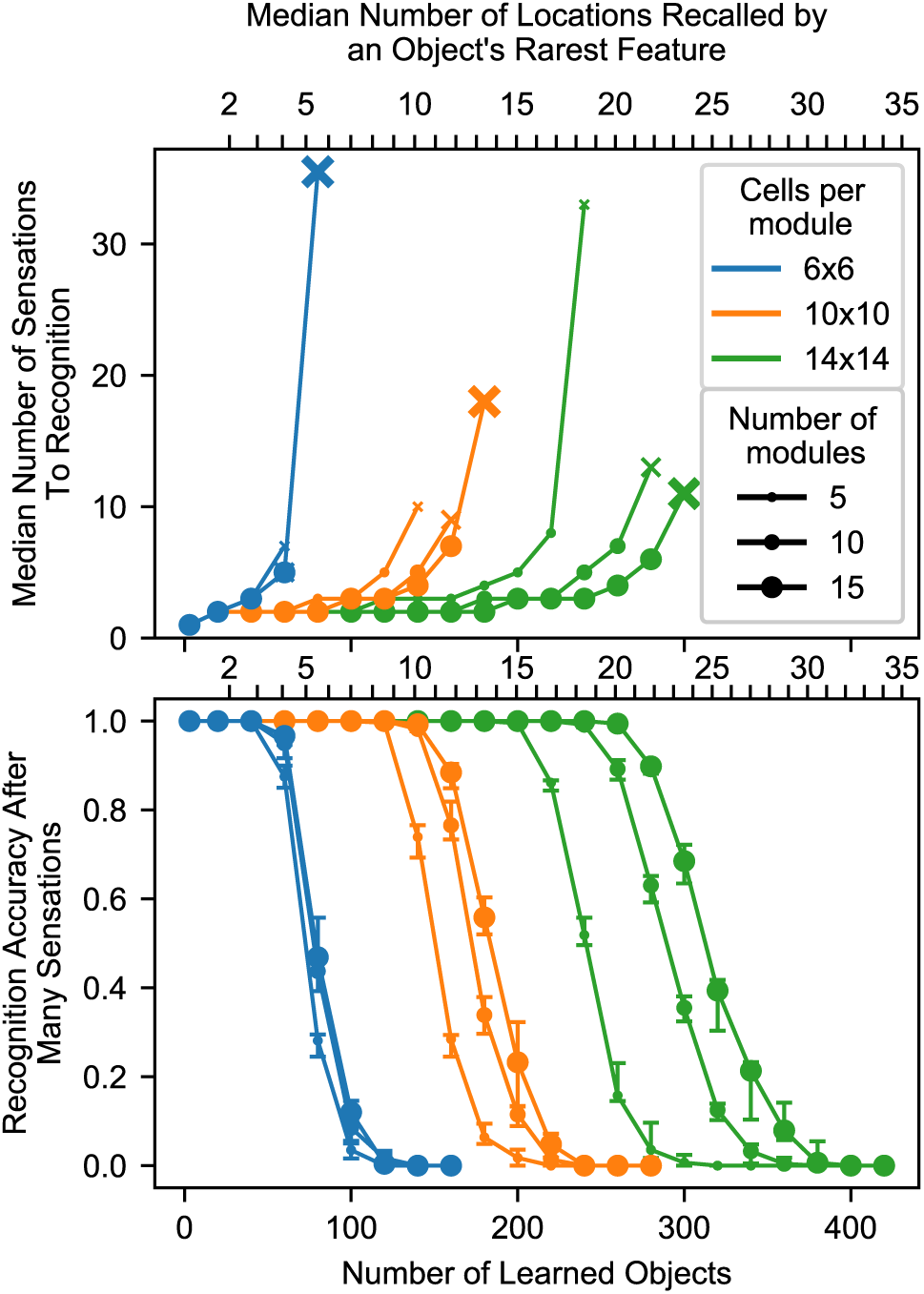
Summary chart showing recognition time and capacity while varying the model parameters. The top chart shows the median number of sensations required to recognize an object, and it marks the final point where a network recognized at least 50% of objects using an ‘x’. The bottom chart shows the percentage of objects that were recognized after traversing every point on the object multiple times. The charts share an x axis, and this x axis has two sets of ticks and labels. The bottom ticks are numbers of learned objects, given 10 locations per object and 100 unique features. The top ticks are the typical smallest union sizes that will be invoked by each object. With different statistics, these curves will shift relative to the bottom ticks, but they should stay approximately fixed relative to the top ticks. We repeat the experiment with 10 different sets of random objects. In the top chart we combine the data from all trials and plot the median. In the bottom chart we plot the 20th, 50th, and 80th percentiles for each data point across trials.

We conclude that this model reliably recognizes an object if the object has at least one sufficiently uncommon feature, a feature that causes the network to recall a sufficiently small number of locations. After sensing this feature, the model can use other, more common features to finish recognizing the object, but it needs some initial clue to help it activate a manageable union that it can then narrow. With enough cells, this initial clue doesn’t need to be especially unique. For example, the feature could have 10 learned locations-on-objects if the module has 10×10 cells per module and 10 modules, as shown in **Figure 9**, and this breaking point can be pushed arbitrarily high by adding more cells. The network can also avoid running into this breaking point by tweaking the set of “features” that it extracts from sensory input.

## MAPPING TO BIOLOGY

We have described a two-layer network model for sensorimotor inference and now consider how this network motif maps to known cortical anatomy and physiology. Layer 4 (L4) is well understood to be the primary target of thalamocortical sensory inputs (Douglas and Martin, 2004; Jones, 1998; Theyel et al., 2010). These connections are believed to be driving inputs (Viaene et al., 2011) and target both excitatory and inhibitory neurons, with a slight delay for the inhibitory neurons that provides a window of opportunity for neurons to fire (Harris and Shepherd, 2015). Mountcastle identified the organization of neurons into mini-columns with shared receptive fields (Buxhoeveden, 2002; Mountcastle, 1957).

The connections between L4 and layer 6a (L6a) closely resemble the connections in our model (**Figure 3**, arrows 2 and 4). Thalamic input forms a relatively small percentage of L4 synapses; approximately 45% of its synapses come from L6a (Binzegger et al., 2004). The connections from L6a are weak (Harris and Shepherd, 2015; Kim et al., 2014). They connect to distal dendritic segments of L4 cells whereas thalamocortical afferents connect more proximally (Ahmed et al., 1994; Binzegger et al., 2004). L4 cells also form a significant number of connections to cells in L6a (Binzegger et al., 2004). These biological details closely match our network model, where the location layer (putatively L6a) has a modulatory influence on the sensory input layer (putatively L4) which in turn can drive representations in the location layer.

Our location layer also requires a motor input. Experiments show that L5 cells in motor regions, such as M2, project to sensory regions, including layer 6 (Leinweber et al., 2017; Nelson et al., 2013). Principal cells in Layer 6 of mouse V1 receive a direct projection from retrosplenial cortex that signal the angular velocity of horizontal rotation of the head (Vélez-Fort et al., 2018). There is also a potential indirect pathway through thalamocortical inputs that target layer 6 (Harris and Shepherd, 2015; Thomson, 2010). Thalamic relay cells receive input from layer 5 neurons that are presumed to be efference copies of motor commands sent subcortically (Chevalier and Deniau, 1990; Jones, 1998). Any of these direct or indirect pathways could serve as motor signals for path integration in L6.

Our model draws inspiration from the grid and place cell systems in the hippocampal formation. Our location layer is modeled after grid cells. These cells project (Zhang et al., 2013) to place cells (O’Keefe and Dostrovsky, 1971) in the hippocampus. Areas of hippocampus containing place cells also project back to areas of entorhinal cortex containing grid cells (Rowland et al., 2013). Many place cells seem to represent item-place pairs, and these pairs are learned through experience (Komorowski et al., 2009), a phenomenon that is analogous to the learning of feature-location pairs in our sensory layer.

Location representations in neocortex similar to grid cells are speculative but there is initial experimental support for grid-like codes in neocortex. fMRI experiments with humans performing tasks have led to activity signatures in prefrontal cortex that are similar to grid cell signals (Constantinescu et al., 2016; Julian et al., 2018a). Direct cell recordings have also shown grid-like activity in frontal cortex (Doeller et al., 2010; Jacobs et al., 2013). These experiments are consistent with the hypothesis that grid-like cells are present in the neocortex and not just a phenomenon of the hippocampal formation. These grid cells were discovered by looking for cells that respond similarly to grid cells in the hippocampal formation; in the Testable Predictions section we predict that more grid-like cells will be found by looking for cells representing sensors’ locations relative to objects.

In our model, primary sensory cortex represents the sensory input at locations in an external reference frame. This is consistent with results from (Saleem et al., 2018). As a mouse ran on a virtual track, the majority of recorded cells in primary visual cortex encoded the animal’s location on the track, even when the mouse received visual input that occurred at multiple points of the track. According to our model, in this task the visual cortex represented the location of a sensor (the mouse’s eye) relative to an object (the virtual track), and the cortex used the visual input to recall locations on the track while using the mouse’s movement to update these location representations. Their finding that the error in V1’s location representation matches the error in CA1’s location representation suggests that these two areas use the same path integration signals.

Although additional experimental work is required, evidence suggests that L4 and L6a provide the best candidate populations for our sensory and location layers, respectively.

## DISCUSSION

We have presented a two-layer neural network model for sensorimotor object recognition. By pairing sensory input with an object-centric representation of location, the model learns objects as spatial arrangements of sensory features. Both object learning and object recognition are independent of the particular order of movements and sensations.

The model’s location layer contains modules that operate similarly to grid cell modules in the medial entorhinal cortex. The location modules represent the sensed location in the reference frame of the object. The location modules receive a movement input that updates and predicts new locations. The sensory layer combines the representation of location with sensory input to create representations of sensory inputs that are unique to objects and locations on those objects.

Object recognition in our model occurs via a series of sensations and movements. A sensory input activates the set of locations where the input has been previously learned. The location layer updates these locations based on motor input. The updated locations cause prediction in the sensory layer. The next sensory input narrows down possible locations and possible objects. In this way, a series of sensations and movements will allow the model to rapidly infer which object is being sensed.

The model provides a concrete implementation of the location signal introduced in our earlier model (Hawkins et al., 2017) and proposes a mechanism for how sensory input and motor input work together.

### Can grid cell modules support unions

In our model we treat a grid cell module as a black box with well-known and previously documented properties. Our model is agnostic regarding the internal dynamics and mechanisms that create the grid cell properties. However, we gave grid cell modules an additional property that is not typically noted in grid cells: support for unions. The grid cell modules in our model can activate, maintain, and shift multiple bumps of activity simultaneously. Various models of grid cell dynamics could be plugged into this model, but this union property introduces a new requirement for these models.

Several models have been proposed to explain grid cell dynamics (Giocomo et al., 2011). Recurrent grid cell models have received more empirical support (Yoon et al., 2013) than models in which grid cells establish their responses independently of each other. In recurrent models, grid cells determine their activity using velocity input and connections to other grid cells, either directly or via interneurons. A well-known recurrent model is the continuous attractor network (Burak and Fiete, 2009; Fuhs and Touretzky, 2006) which performs robust path integration and offers a simple explanation for the origin of the hexagonal firing fields. Another explanation for the origin of these fields is that they are an optimal code for locations which is naturally learned by neural learning rules. This argument has appeared in two lines of research. In path integration models, (Banino et al., 2018) and (Cueva and Wei, 2018) found that recurrent neural networks trained to perform path integration naturally develop grid cells, although neither report the network developing the full rhombus of grid cells at each scale. Setting path integration aside, (Kropff and Treves, 2008), (Dordek et al., 2016), and (Stachenfeld et al., 2017) showed that cells performing Hebbian learning on place cell activity would naturally learn periodic firing fields similar to those of grid cells.

The continuous attractor model is so named because it has a continuous manifold of stable states. If the network activates a representation that isn’t within this manifold of stable representations, the activity will move to the nearest stable state. In typical continuous attractor networks, a union is not a stable state and the network will collapse a union of bumps into a single bump. It’s an open question whether it’s possible to have a continuous attractor with stable union states. In this paper we’ve shown that it’s theoretically advantageous for grid cell modules to work with unions. The attractor dynamics are appealing in part because they explain the hexagonal firing fields, but as mentioned, those may be explainable as the natural result of a recurrent neural network learning a location code.

Modeling the internal mechanisms of the location layer is an area for future research.

### Egocentric vs. object-centric coordinates

Sensors such as eyes and skin detect features in a viewer-centric or egocentric coordinate frame. It is inefficient to learn an object’s features in egocentric coordinates as the system will need to learn the object at every shifted location and rotated orientation. Our model represents location using grid cell-like modules, and, like grid cells in the entorhinal cortex, cortical grid cells represent locations relative to the external object being observed.

Converting from an egocentric reference frame to an object-centric reference frame is therefore necessary. If reference frames are based on Cartesian coordinates then it is necessary to establish origin points and the conversion, as outlined in (Marr and Nishihara, 1978), is complex. Grid cell representations avoid much of this complexity. Reference frames based on grid cells do not have an origin. Grid cells represent locations and features relative to each other as opposed to relative to an origin.

However, representing objects with grid cells still requires knowing the orientation of the sensor and features relative to the object. Our model does not yet have a representation of orientation. As a result, our model will only recognize an object if the object is at its learned orientation relative to the sensor. An extended version of our model could incorporate orientation using analogs to head-direction cells (Taube et al., 1990). Grid cells represent the animal’s location on a cognitive map, whereas head-direction cells represent the animal’s orientation relative to the cognitive map. When an animal moves, grid cell modules move their bump of activity depending on the orientation of the animal. Similarly, we expect the neocortex to represent the orientation of sensors relative to the reference frame of the sensed object, and this orientation will influence how sensor movement translates into the movement of bumps in cortical grid cell modules. This extended model would be able to learn orientation-invariant models of objects as well as handle sensors that rotate with respect to objects.

### 2D vs. 3D objects

We have described our model using 2D grid cell modules to learn 2D objects, however, the neocortex is capable of learning 3D objects. Our model should work with 3D objects provided the location code represents 3D locations relative to an object. How the entorhinal cortex represents 3D space is an active area of research (Jeffery et al., 2015). Our team is currently working on extending our model to include 3D representations and orientation.

### Relationship to other models

Our model identifies objects using the relative location of sensory features. Objects are disambiguated over time through successive sensations and movements. In contrast, most existing models of object recognition involve a strictly feedforward spatial hierarchical system (DiCarlo et al., 2012; Riesenhuber and Poggio, 1999; Serre et al., 2007; Yau et al., 2009). In these models each level detects the presence of increasingly abstract features in parallel until a complete object is recognized at the top of the hierarchy. Our model implies that each level of a hierarchy might be more powerful than previously assumed. In (Hawkins et al., 2017) we discussed how spatially separated sensory inputs (across multiple cortical columns each computing a location signal) can cooperate in parallel to recognize objects, and some of the implications on hierarchy. Our model suggests a path for integrating sensorimotor behavior into a hierarchical system, and accounts for the many inter and intracortical connections that are not explained by a purely feedforward model. A more detailed study integrating our model into a full hierarchical system is a topic for future research.

There is significant literature on sensorimotor integration and the learning of internal models in the context of skilled motor behavior (Wolpert et al., 2011; Wolpert and Ghahramani, 2000). These have primarily focused on learning motor dynamics and kinematic control, including reaching and grasping tasks. Our model focuses on the more structured object recognition paradigm, but there are many high-level similarities with this body of literature. Our location layer is highly analogous to the forward models posited to exist in motor control (Wolpert et al., 2011). In both cases the current state is updated using a motor efference copy to compute the next state. In both these models, this is an estimate of the state that informs predictions (arrow 2 in **Figure 3**) and is then combined with sensory input to produce the current state (arrows 3 and 4 in **Figure 3**). The primary difference is that our model recognizes a set of structured objects rather than motion trajectories. The neural mechanisms are also significantly different. Nevertheless it is intriguing that the same ideas can be applied to both situations and may reflect a more general design pattern in the brain. An in-depth exploration of this relationship is a topic for future research.

Our model provides an alternate explanation for predictive processing in visual cortex. Existing models of saccadic remapping suggest that it occurs by shifting attended parts of the image across visual cortex (Wurtz, 2008). This explanation requires every part of the retinotopic map to be connected to every other part, either horizontally or through the feedforward input, and it requires using the eye movement information to enable a small subset of these connections. In our model, each patch of visual cortex computes the location of a patch of retina relative to the attended object, then uses this location to predict sensory input. As the eyes saccade over a static object, our model would not require any horizontal shifting of information within the visual cortex. Thus this paper suggests a model of saccadic remapping that does not require every patch of visual cortex to be related to every other patch via long distance connections. Implementing this extended model is a topic for future research.

A recent article (Keller and Mrsic-Flogel, 2018) proposed a neural circuit for using a signal to form predictions and represent prediction error. This is a general-purpose circuit that can consume any type of prediction signal. In the present study we have presented a particular type of prediction signal – locations – and how they are updated. We combined this signal with a different neural mechanism for prediction (Hawkins and Ahmad, 2016), but it would also be possible to use locations as prediction signal with Keller and Mrsic-Flogel’s mechanism. The two mechanisms differ in how they represent prediction error. In our model, when an input matches a prediction, the network activates a sparse representation of the input in this particular context, whereas a mismatch causes the network to activate a dense representation of the input in many different possible contexts. Keller and Mrsic-Flogel’s model represents sensory stimuli and prediction error using two different populations of cells.

Others have emphasized the importance of sensorimotor processing in how we perceive different sensory modalities differently. (O’Regan and Noë, 2001) use the example of holding a bottle and seeing a bottle. They propose that your conscious perception of the bottle via a sensory modality comes from your mastery of that modality, from being able to predict what you will feel as you move your hand over the bottle or what you will see as you move your eyes over the bottle. In this paper we have not focused on conscious perception, but our model does propose how the brain represents the bottle in different sensory modalities in such a way that it will make predictions in response to movements. We think the idea of object-specific location representations is quite compatible with this view of perception.

A recent article from our lab (Hawkins et al., 2019) proposed a location-based framework for understanding the neocortex. There we described how models of objects can be related to one another by representing displacements between grid cells, enabling representations of compositional objects. In the present paper we provide a neural mechanism for learning object models that fit into this approach to compositionality. This is one building block of a larger theory in which locations are key computational primitives of the neocortex.

### Testable Predictions

Our model makes a number of experimentally testable predictions. We expand on the predictions from (Hawkins et al., 2017).

1. The neocortex uses analogs of grid cell modules to represent locations relative to objects. The cell activity in a module moves through a manifold of representations as the attended location moves relative to an object. For example, in somatosensory areas, cells will respond selectively when the animal’s finger is at particular locations relative to an attended object. Just as entorhinal grid cell modules use the same map for every environment, the cells of a single module use the same manifold of representations for every object. This map has limited size, and hence it will perform some form of wrapping at its edges.
2. The neocortex uses a population code of multiple modules to represent object-specific locations.
3. These modules are in Layer 6 of the neocortex.
4. The projection from Layer 6 to Layer 4 modulates which cells in Layer 4 become active. If Layer 6 input is experimentally inhibited, activity in Layer 4 will become denser.
5. The connection from Layer 4 to Layer 6 can drive the Layer 6 cells to become active, but this only occurs when the animal receives an unpredicted input.

## MODEL DETAILS

Each module has a fixed number of cells which each have a fixed phase in the rhombus. Gaussian bumps of activity move over these cells. A cell is considered active if its firing rate is sufficiently high. In this section, we walk through the details of these calculations.

Each module contains *w* * *w* cells. Each cell *c* has a constant phase 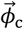 We partition the 2D range [0,1) × [0,1) into *w* * *w* ranges of equal area and set each cell’s 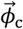. to the center of one of these ranges. Because these modules use basis vectors separated by 60° (Eq. (3)), when mapped onto physical space these cells form a rhombus and they pack together in a hexagonal formation.

The normalized firing rate of a cell *c* caused by bump *b*, denoted *r*_*c,b*_, is equal to the Gaussian of the distance between them.

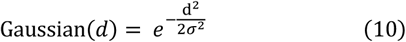

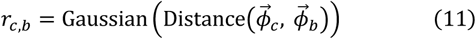

Distance 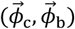 represents the shortest distance between the cell and the bump on the phase rhombus. Computing this distance requires changing the basis so that each 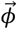 is a point on a rhombus rather than a point on the [0,1) × [0,1) square. *σ* specifies the size of the bump relative to the rhombus, which we discuss later.

When there are multiple bumps, the firing rates from each bump are combined as if each rate encodes a probability of an event. The combined firing rate encodes a probability of the “or” of those events.

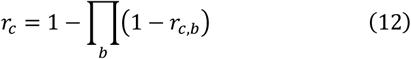

To compute module *i*’s active cells 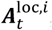, we compute each cell’s firing rate and check whether it is above the active firing rate *r*_active._

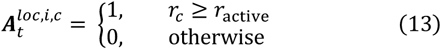

We choose *r*_active_ using a readout resolution parameter *δϕ* which is common in grid cell models (Fiete et al., 2008; Sreenivasan and Fiete, 2011).

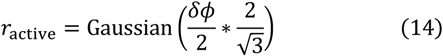

The readout resolution *δϕ* approximately specifies the diameter of the range of phases that a bump encodes. Because modules are 2D, if the readout resolution is 1/4 then the bump can encode approximately 16 possible positions in the rhombus. The multiplicative factor of 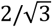 accounts for the fact that when circles pack together in hexagonal formation, they leave some area uncovered; this factor expands the circles to overlap and cover this area. With a single bump, changing the *σ* parameter has no effect on the model, because the *δϕ* parameter has complete control over what fraction of the cells are considered active. *σ* becomes relevant when there are multiple bumps. The wider these bumps are, the more they’ll combine and cause interstitial cells’ firing rates to rise above *r*_active._

We vary *σ* and *δϕ* as follows. Our baseline parameters mimic the sparsity of rat entorhinal grid cell modules. We set *σ* to 0.18172, a number we obtained by fitting a 2D Gaussian to the firing fields generated by the model in (Monaco and Abbott, 2011). We set *δϕ* to a conservative estimate of 1/3. In this configuration, we assign the network 6×6 cells, and a bump always activates at least 2×2 cells. When we test the network with more cells, we assume that the bump remains a fixed size relative to the cells, i.e. that the bump is smaller relative to the size of the module, and we scale down *σ* and *δϕ* accordingly. For example, with 12×12 cells, we use *σ*/2 and *δϕ* / 2. By varying the parameters in this way, a single bump always activates between 4 and 7 cells, depending on where the bump is centered relative its local neighborhood of cells, and as we vary the number of cells we’re effectively varying the number of cells that the bump *doesn’t* activate.

During learning, only the cell with the highest firing rate is associated with the sensed feature. (When a sensed feature activates a cell, it activates a bump centered on the cell via Eq. (7), activating the cells around it.) This means this model’s learning resolution is twice as precise as the readout resolution. Because sensed features are associated with cells that represent a range of phases, there’s always some uncertainty in the phase recalled by sensory input. If the learned resolution weren’t more precise than the readout resolution, the bump of active cells would need to expand to account for this uncertainty, and the effective readout resolution would be half as precise. Using fixed-sized bumps, achieving a particular readout resolution – that is, having a bump encode a range of phases with a particular diameter – requires the learning resolution to be at least twice as precise as this readout resolution.

The ideal classifier in **Figure 7B** stores all objects as 2D arrays. During inference, it uses the first sensed feature to find all possible locations on all objects with that feature, and it stores these as candidate locations. With each subsequent movement it updates all of the candidate locations. Any updated candidates that are not valid locations on objects or contain features that don’t match the new sensed feature are removed from the candidate list. Once there is only a single location left, inference is successfully completed.

The bag of features detector stores a set of features for each learned object. It does not keep track of how many times features occur, just the set of unique features present somewhere on the object. During inference, another set keeps track of which features have been sensed so far. Once there is only one object that contains all of the sensed features, inference is successfully completed. If there are multiple objects that contain all features once all locations on the object being tested have been visited, then the object cannot be uniquely classified.

### Recurrent dynamics

This network contains a recurrent loop, so there is potential for additional recurrent dynamics after Stage 4 of a given timestep. However, because the input in Stage 2 is modulatory and never drives cells to become active, the network generally converges after Stage 4. We found that simulating these dynamics occasionally allowed the network to recognize objects with one fewer sensation. To keep the model and the notation simple, in this paper after Stage 4 the simulation advances to the next timestep, and Stage 1 repeats with the next sensory input.

### Code availability

All of the source code for this model and these simulations can be found at https://github.com/numenta/htmpapers.

**Figure S1.**
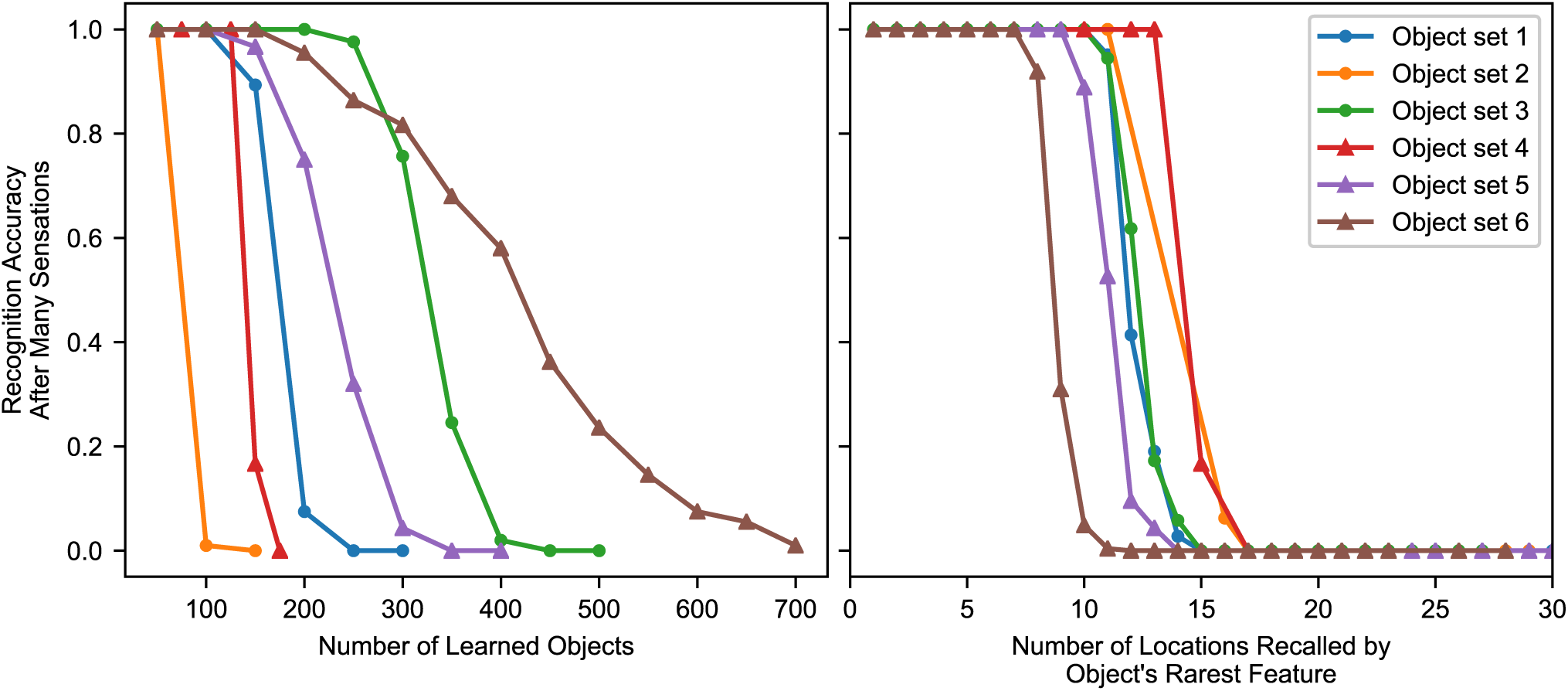
Varying the object statistics, the model’s breaking point varies significantly relative to number of learned objects. The breaking point is much more consistent relative to the number of locations recalled by object features. In these charts we use a single model and test on 6 different distributions of objects. The model uses 10 modules with 10×10 cells per module. **(Left)** The network’s capacity depends on the statistics of objects. The network’s performance begins to break down after a certain number of objects, and this breaking point can vary by orders of magnitude with different object distributions. **(Right)** This breaking point varies significantly less when described in terms of “number of locations recalled by a sensation” rather than “number of objects learned”. Using the same data from the first chart, for each object we measure the total number of occurrences of the object’s rarest feature, and we plot recognition accuracy against this number. With each of these object distributions, the model reaches its breaking point when the number of recalled locations is within a small interval – conservatively, between 7 and 15. There is still some variation due to the statistics of the object’s other features (not just its rarest feature), but the number of occurrences of the rarest feature provides a good first approximation for whether the network will recognize the object. **(Object descriptions)** Each object set had 100 unique features and 10 features per object, except where otherwise noted. The first three sets generate objects using the same strategy as all the other simulations, varying the parameters. The last three use different strategies. *Object Set 1:* baseline. *Object Set 2:* 40 unique features rather than 100. *Object Set 3:* 5 features per object rather than 10. *Object Set 4:* Every feature occurs the same number of times, +/-1, rather than each object being randomly selected set of features with replacement. *Object Set 5:* Bimodal distribution of features, probabilistic. Divide features into two equal-sized pools, choose features from the second pool more often than features from the first. *Object Set 6:* Bimodal distribution of features, enforced structure. The features are divided equally into pools. Each object consists of one feature from the first pool and nine from the second.

